# Reliable Identification of Homodimers Using AlphaFold

**DOI:** 10.1101/2025.11.27.691011

**Authors:** Sarah Narrowe Danielsson, Arne Elofsson

## Abstract

**Motivation:** Protein-protein interactions are central for understanding biological processes. The ability to predict interaction partners is extremely valuable for avoiding costly, time-consuming experiments. It has been shown that AlphaFold has an unsurpassed ability to accurately evaluate interacting protein pairs. However, a protein can also form homomeric interactions, i.e. interact with itself.

**Results:** We found that AlphaFold yielded a significantly higher false-positive rate for identifying homodimers than for heterodimers. True Positive Rate (TPR) at 1% False Positive Rate (FPR) drops from 63% for heterodimers to 18% for homodimers. When we investigated the high-scoring false positives, i.e., non-homodimers with high AlphaFold scores when predicted as such, we found that their homologs were enriched for homomultimeric proteins. Using a simple logistic regression model that combines AlphaFold scores with structural and homology information, we increased the TPR (at 1% FPR) to 42±8% (5-fold cross-validation) from 19%. If we excluded the homology information, we achieved a TPR of 28±7%, which is still better than using AlphaFold metrics.

**Availability and implementation:** All data are available from Zenodo DOI:10.5281/zenodo.17738668 and all code from https://github.com/SarahND97/alphafold-homodimers.

**Supplementary information:** Supplementary information is available online.

## Introduction

Protein-protein interactions are central to the understanding of biological processes. However, uncovering new interactions is challenging due to their diversity, which ranges from long-term, stable complexes to short, unstable, transient interactions. In addition, a protein chain can have multiple interaction partners and form self-interactions. All of these factors combined make the problem challenging both experimentally and computationally. Also, not only is it important to know which proteins interact, but it is crucial to understand how they interact, e.g., by determining their 3D structures (1).

Determining the structure experimentally is expensive, time-consuming, and near impossible for certain classes of proteins. It is therefore useful to be able to predict the structure of a protein computationally from its amino acid sequence. In 2021, DeepMind released AlphaFold2.0 (AF2) (2), which was able to predict monomers with experimental precision and, with some tweaks, also multimers (3; 4; 5; 6; 7; 8). In 2022, AlphaFold-Multimer was introduced, a version of AlphaFold2 trained on multimers with known stoichiometry (9). The latest version of AlphaFold-Multimer (AlphaFold2.3) was released in 2022 (10).

Recently, AlphaFold3 (AF3) was released(11). In contrast to AF2, AF3 can predict all types of molecules present in the Protein Data Bank. The inference source code was released in November 2024, enabling large-scale usage (12).

When using AlphaFold to predict protein-protein interactions, it is important to note that all versions primarily predict a structure rather than an interaction score. This means that we have to infer an interaction score from the quality of the predicted structure. When Bryant et al first used AlphaFold2.0 for interaction prediction, we developed our own score pDockQ for this purpose (3). After the release of AlphaFold-Multimer, it is possible to infer this from the quality metrics produced by AlphaFold, primarily the interface-predicted TM-score (ipTM) and the ranking confidence, which is a combined score of ipTM and pTM.

Bryant et al showed that AlphaFold2.0 could be used to successfully separate interacting and non-interacting heterodimers (3). Additionally, Schweke et al. showed that by using the predicted aligned error (pAE) and information about contacts in the predicted structure, AlphaFold2.0 could be used to predict whether a protein forms a biologically feasible homodimer (13). In their study, they used ColabFold, which made AlphaFold2.0 (both for monomers and multimers) available in a Google Colab notebook in 2021 and has since been extended to include AlphaFold2.3 and Deepfold (5; 14).

The main difference for AlphaFold when predicting homodimers and heterodimers is that homodimeric interactions do not require any pairing in the multiple sequence alignment (MSA), making them “easier” problems for pure complex prediction. However, that could also make it more difficult to separate proteins that form homodimers from those that do not.

In this study, we investigated differences in separating interacting and non-interacting protein pairs, when the chains are identical (homodimers) and when they are different (heterodimers). We found that AlphaFold has a lower true-positive rate (TPR) at low false-positive rates (1% and 5%) for homodimers than heterodimers. We sought to understand why and found that the answer might lie in the stoichiometry of homologs. We observed that stoichiometry varies across all protein classes. Moreover, proteins that have many homologs of a different stoichiometry are more likely to be wrongly predicted by AlphaFold. We combined homology and interface information with AlphaFold confidence metrics and found that we could improve the TPR at a 1% FPR from 19% to 42±8%. The most important feature was the homology information, without which the TPR was only 28±7%. This is still significantly lower than for heteromeric interactions (TPR = 63%), indicating that the problem is not fully resolved.

## Methods

### Dataset Generation

The proteins used for the dataset were taken from the Protein Data Bank (PDB) on the 10th of March 2025 (15). All of the proteins were released after September 30th 2021, since this is the latest training cut-off used by any AlphaFold version. We collected proteins from three categories: homodimers, heterodimers, and monomers. All individual chains in each category were then clustered using MMseqs2(16) using a minimum sequence identity of 30%, and the centerpoints of each cluster were selected. This was to avoid bias towards a certain class in the test set.

The proteins were filtered to ensure they did not contain any DNA/RNA, had a resolution of 4.5 Å or better, and NMR structures were excluded. We also used only entries containing exactly one bio-assembly to avoid cases where different bio-assemblies of the same PDB entry have different stoichiometries. After careful consideration, we decided not to compare our selected proteins to the AlphaFold training set. Removing all homologs between our test set and the AlphaFold training set would significantly reduce the size of the test set. Therefore, we decided to use only the time split. This would also represent a real-world scenario. The data selection process is visualised in Figure 1.

**Fig. 1.**
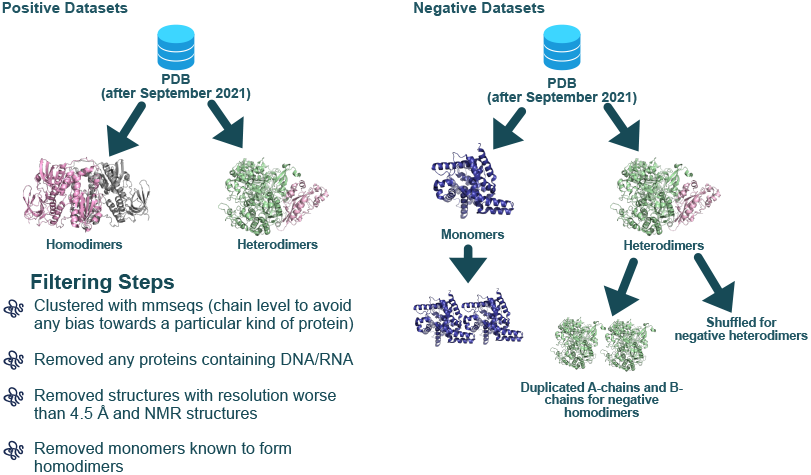
An overview of the dataset generation process.

We also ensured that the monomeric dataset did not contain any sequences found in multimers. We did this by using MMseqs2 easy-cluster with a 90% sequence identity cut-off on the entire PDB, and then removed any monomers in our dataset that were present in clusters that contained multimers. Some proteins from all datasets were also excluded because their sequences were too long to be predicted by one of the three AlphaFold versions used in the study. The PDB Ids used in the study are available on Github: https://github.com/SarahND97/alphafold-homodimers. In the end, our dataset consisted of 288 monomers, 169 heterodimers, and 466 homodimers, for a total of 922 homodimeric pairs. For the heterodimers, we also created a negative dataset by shuffling chains and ensured that no pair remained identical. This gave us a total of 146 shuffled heterodimers and 146 positive heterodimers.

### AlphaFold Settings

In this study, we assessed the performance of AlphaFold2.0, AlphaFold2.3, and AlphaFold3. The multiple sequence alignments (MSAs) used by all AlphaFold versions were created with MMseqs(16) using the default settings of ColabFold(5). We also tried altering the MSA settings, specifically the E value, to see if this improved performance. All methods were run with default settings, without templates, and 25 structures were predicted in total (5 per AlphaFold model). The exception was ColabFold, where 5 structures were predicted, as the default for AlphaFold2.0 was to generate 5 models, which was later extended to 25 for AlphaFold2.3 and AlphaFold3.0. From these 25 (or 5) models we also calculated the minimum, maximum and average of the confidence metrics produced by AlphaFold.

We tested running AlphaFold 2.0 with both the FoldDock and ColabFold pipelines. After a quick inspection, we noted that the ColabFold pipeline performed significantly better for homodimers, producing fewer instances of complete chain overlap. We assume this is due to ColabFold not pairing the MSA for homo-oligomers, instead using gapped alignments, as the original ColabFold authors stated that this improved their results (5). Therefore, FoldDock was ignored for further analysis. The main reason for including ColabFold (5) was to investigate the QSproteome logistic regression model used in (13), which is based on predictions of 5 structures.

### Identification of Interacting and Non-Interacting Pairs

The quality of an AlphaFold model can be estimated from internal ipTM or ranking confidence values, as well as from external methods that evaluate the model’s structure independently or in combination with internal AlphaFold metrics, such as pLDDT. Further, we can use these values for a single model or for a combination of all (25) models, such as the minimum, maximum, or average of the ipTM. In this study, we focus on the ranking confidence since it is common practice to use the highest-ranked structure. The ranking confidence (for complexes) is a combination of 80% ipTM and 20% pTM, indicating that ipTM and the ranking confidence correlate strongly.

### Conservation of Stoichiometry of Homologs

We used Foldseek (17) to identify similarities between the structures in our dataset and those in the PDB (updated on the 26th of June 2025). We assumed that most of the hits we found were homologs of proteins in our dataset, especially at high TM scores; at low TM scores, these might show similarity due to convergent evolution. We compared both the predicted and native structures to the PDB and observed minimal differences, as expected, given the high accuracy of the predicted structures. Therefore, we decided to use the native structures. In addition, we used MMseqs2 to search the same set of proteins.

We classified all detected homologs from Foldseek (and MMseqs2) into three main stoichiometry categories: monomers, homomultimers, and heteromultimers, based on the chains of the biological units. Next, we computed the fraction of each category at different TM score cut-offs.

### Features for Homodimer Prediction Improvement

Given the subpar ability of AlphaFold to identify homodimers, we set out to improve this. The main target function we aimed to improve was the TPR at 1% FPR, using additional information about protein pairs. Inspired by the SPOC model (18), we extracted features from the predicted complexes and their interfaces. In addition, we used various versions of the AlphaFold confidence metrics, along with stoichiometry information among homologs. We used both information from a single AlphaFold model and from the set of 25 models generated for each pair. All these values were tested in various combinations using a simple logistic regression model to find the best TPR.

#### Homology Information

From the homology data, we removed any Foldseek matches with *Fident* above 0.5 (50% sequence identity) to the query using MMseqs2. From the remaining Foldseek hits, we derived the fractions of homomultimers (*HM Frac*) and the fraction of all (homo+hetero) multimers (*Multimer Frac*) at different TM-score cut-offs. When there were no hits, the fractions were set to zero. We also retrieved the highest TM score for homomultimers, multimers and all matches. In total, we ended up with 23 features summarizing the stoichiometries of the homologs in our dataset, see Supplementary Table S1.

#### AlphaFold Confidence Metrics and Structural Consensus

From AlphaFold, we gathered the ipTM and ranking confidence scores from the 25 predicted structures. We calculated the minimum, maximum and average for each of these scores.

We have noted that often when a prediction is correct by AlphaFold, all 25 models are very similar, but when they are wrong, they are often different (19). Therefore, we also used USAlign(20) (with the flag -mm 1) to calculate the average similarity between all 25 predicted structures of one target (excluding self-comparisons and symmetric comparisons), resulting in 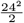 comparisons. The average TM score will be referred to as the *Structural Consensus*.

#### Properties of Interface Surface

(18) used several different ways to characterise contacts and pairs of residues in the interface of a predicted protein-protein structure to develop a method (SPOC), aimed at improving the identification of human-interacting heteromeric protein pairs. In the original SPOC model, 42 features were initially included, which were later pruned to 24. Of these 24, there were 12 structural features. Here, we used the same 12 structural features to improve homodimer identification. SPOC also uses genetic database information and AlphaMissense features, but because these are only available for human proteins, we decided to exclude them.

The final 12 pruned structural features include *min contacts across predictions, num contacts with max n models, num unique contacts, mean contacts across predictions, best num residue contacts, best if residues, best plddt max, best contact score max* and *Best pAE min*. These 12 features also include the minimum, maximum, and average ipTM values as mentioned in Section 2.5.2. For an explanation of each feature, see Table S2 in the Supplementary Material and (18).

Next, to analyse the interface surface, we used the tool FreeSASA(21) to calculate the Solvent Accessible Surface Area (SASA). We extracted the buried a/polar size of the interface and used both the sizes and the a/polar fraction. Here, we calculated these metrics for each of the 25 predicted models and then used the average as the final result.

### Feature Clustering

In total, we have 44 features (15 interface/structural features, 23 homology-based, and 6 from AlphaFold) that can be combined to develop a logistic regression-based predictor. Testing all combinations of these 44 features would have been ideal. However, this would have required us to try 44! combinations, which was simply not feasible. Therefore, we decided to reduce the number of features by using hierarchical clustering.

Before clustering, missing values were imputed, near-constant features were removed, and the remaining features were normalized. We then defined feature similarity as the absolute Spearman correlation, transformed it into a distance measure (1 – correlation), and applied average-linkage hierarchical clustering. We first clustered the homology information features into 6 clusters, selected the best-performing features from each, and then clustered all 29 features into 11 clusters. See Figure S5 in the Supplementary Material for final clusters.

### Fitting a Logistic Regression Function to the Features

We used a 5-fold cross-validation scheme to find the optimal set of features for the logistic regression function. The primary model used in the study was optimized for the best TPR at a 1% FPR. We tried all combinations using 0 or 1 features from each cluster. We also tried excluding homology information to test its effect on the performance of the function. This was important to test, as there are proteins with little to no homology to other proteins. We saw that this significantly reduced performance, as shown in detail in Table 4 and Figure 4. We reported the results as the average across the 5 folds, along with the standard error of the mean. For the released logistic regression functions, we used the entire dataset.

## Results and Discussion

It is well-known that AlphaFold is very good at separating interacting from non-interacting heterodimeric protein pairs (22). A large fraction of all interacting pairs have high ipTM/ranking confidence scores, whereas this is extremely rare for two randomly chosen proteins (3). However, to the best of our knowledge, it is not well established whether the same holds true for homodimers; i.e., whether proteins that interact with themselves achieve a high ipTM/ranking confidence score, and proteins that do not form homodimers achieve low ipTM/ranking confidence.

### AlphaFold Produces High Scoring False Positives for Homodimers

First, we assessed AlphaFold’s ability to separate interacting and non-interacting protein pairs in our test set. A major challenge in this study was defining a reliable negative dataset for homodimer prediction, as it is difficult to confirm that a protein does not form a homodimer, even under high concentration.

We inferred the stoichiometry from the PDB bio-assembly .*cif* files as our ground truths, acknowledging that they may contain errors. To minimise the effect of potential classification errors in the negative dataset, we used two different sets. The first dataset consisted of monomers and the second of separated heterodimeric chains (see Section 2.1). The results were similar for both negative datasets, with only a slightly higher TPR at 1% FPR for monomers, while the heterodimer-based negatives showed a slightly better area under the precision-recall curve (AUPR) (the Table 1).

**Table 1.** Area under the precision-recall curve (AUPR), F1-score and MCC-score. F1-score and MCC-score are calculated using the highest ranking confidence for each PDBid, and the threshold used for the calculation of F1 and MCC is 0.8. The best value for each metric is marked in bold.

| Dataset | AUPR (↑) | F1 (↑) | MCC (↑) | TPR@5%FPR (↑) | TPR@1%FPR (↑) |
| --- | --- | --- | --- | --- | --- |
| Homodimers, Neg: Monomers | 0.91 | 0.81 | 0.59 | 0.52 | 0.22 |
| Homodimers, Neg: Heterodimers | **0.94** | **0.83** | 0.54 | 0.52 | 0.16 |
| Homodimers Both Negative Datasets | 0.86 | 0.79 | 0.61 | 0.52 | 0.18 |
| Heterodimers | 0.91 | 0.77 | **0.67** | **0.69** | **0.63** |

Examining the precision-recall curves 2A and the area under the precision-recall curve (AUPR) in Table 1, it seems that AlphaFold performs similarly for positive homodimers and heterodimers. However, in contrast to heterodimers, only about one-third as many (18% vs. 63%) of the positive set can be detected at an FPR of 1%. This can be explained by a significantly higher number of high-scoring negatives, as seeb in Figure 2B and C. In the density plot in Figure 2B, the main difference in performance is that both homodimeric negative sets produce predictions with ranking confidence above 0.8, while these are almost nonexistent for the shuffled heterodimers. This trend can be seen for all versions of AlphaFold and QSProteome. Therefore, for the remaining part of this study, we focus on the results of AlphaFold2.3; while for the results of the other AlphaFold versions and QSProteome we refer to the Supplementary Material Table S1.

**Fig. 2.**
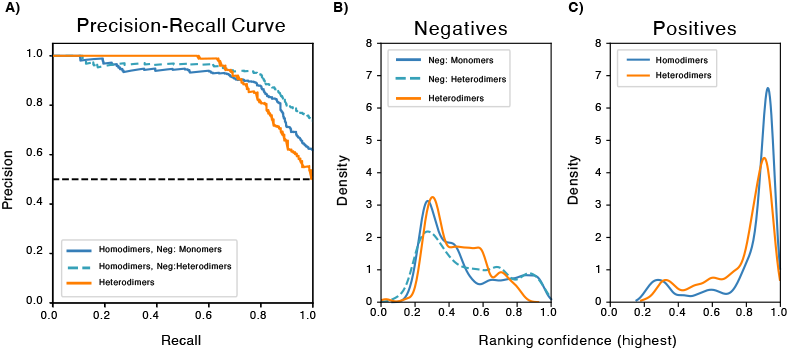
The results of AlphaFold predicting homodimerization vs heterodimerization. A) Precision-recall curves. We can clearly see that the precision remains close to 1.0 for recall between recall 0.0 and ≈ 0.6 for heterodimers, while for homodimers the precision drops already around recall 0.1. Despite this observation, the final AUPR is higher for homodimers. B) Density plots for the negative datasets. We can see that for heterodimers, there are no negatives with a ranking confidence above 0.8. For the two homodimeric negative datasets, we have several predictions with ranking confidence above 0.8. C) Density plots for the positive datasets. Here, we see that the performance is similar to heterodimer prediction, yielding a slightly larger proportion of positives with a ranking confidence of 0.6.

#### Difference Between Prediction of Homo and Hetero-dimers

Next, we sought to find the underlying reasons for the high-scoring negatives. There are two fundamental differences between predicting hetero- and homo-dimeric interactions from any method, such as AlphaFold, based on co-evolution. First, to predict heterodimeric interactions, it is necessary to “pair” the two multiple sequence alignments to identify inter-chain co-evolutionary signals; this is not necessary for homodimers, as they interact with themselves. This should improve the prediction of homodimers, as pairing often results in the information from unpaired paralogs being ignored. Secondly, in a homodimeric interaction, co-evolution occurs for both inter- and intra-chain contacts, and separating these might require structural information of the monomer (23). However, given the highly accurate predictions of AlphaFold, structural information of the monomers should exist for most pairs.

AlphaFold uses multiple sequence alignments to predict protein structures. Therefore, one possible reason for the high false-positive rate might be the presence of homodimers (or homomultimers) within the MSA. To examine whether a stricter cutoff or focusing on orthologs could decrease the number of false positives, we regenerated the MSAs using stricter E-value cutoffs and only 1 iteration or by using only the top-scoring homolog in each species; see Table 2. We may see a small (not significant) improvement by changing the default E-value from 10^−1^ to 10^−4^, but using even stricter E-values actually decreased performance. This indicates that either the change in stoichiometry occurs even among close homologs or that there is some other reason for the high-scoring false positives.

**Table 2.** Area under the precision-recall curve (AUPR), F1-score and MCC-score for the different MSA settings. The thresholds for F1 and MCC are 0.8. All measured targets are scored by the maximum ranking confidence across all 25 models generated.

| MSA Setting | AUPR (↑) | F1 (↑) | MCC (↑) | TPR@5%FPR (↑) | TPR@1%FPR (↑) |
| --- | --- | --- | --- | --- | --- |
| E-Value==$10^{-1}$ (Default) | 0.86 | 0.79 | 0.61 | 0.52 | 0.17 |
| E-Value==$10^{-4}$ | 0.86 | 0.79 | 0.61 | 0.51 | 0.20 |
| E-Value==$10^{-11}$ | 0.86 | 0.79 | 0.62 | 0.50 | 0.15 |
| E-Value==$10^{-60}$ | 0.82 | 0.71 | 0.53 | 0.48 | 0.20 |
| Orthologs (1 Seq Per TaxID) | 0.86 | 0.79 | 0.61 | 0.51 | 0.13 |

#### How Conserved is Stoichiometry Among Homodimers In Our Dataset?

Before diving into the results of our predictions, we examined how conserved the stoichiometry of a protein is in PDB. We used Foldseek to find homologous proteins as described in Section 2.4. We define three classes of stoichiometry; Monomers, Homomultimers and Heteromultimers. Consequently, in Figure 3A-C, we measured the fraction of Monomers, Homomultimers and Heteromultimers for all Foldseek matches of our proteins in the three homodimeric datasets at different TM-score cutoffs. Figure 3 shows that all three datasets have homologs belonging to all categories at all TM-score cutoffs below 0.9.

**Fig. 3.**
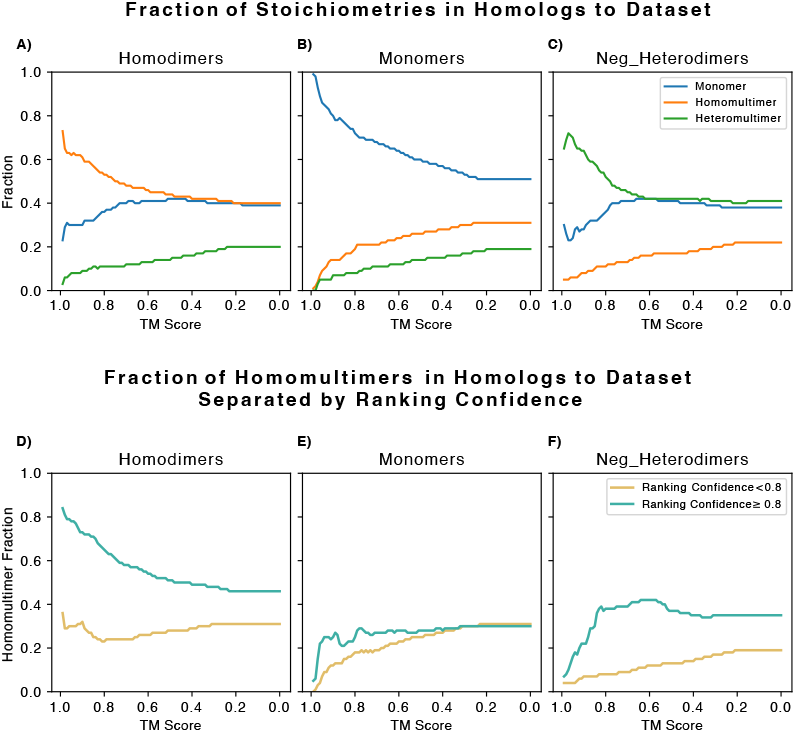
Fractions of stoichiometres of homologs at different TM score cut-offs. A)-C) The stoichiometry fractions of the three datasets, homodimers, monomers and negative heterodimers. The three stoichiometry fractions were Monomer, Homomultimer and Heteromultimer. We can see that in all three datasets, there are similar structures for all stochiometries. D)-Focusing on only the homomultimer fraction here for the homologs of the three datasets. Across all three datasets, we see that mispredicted proteins have a higher/lower fraction of homomultimers.

In more detail, we can see that for Homodimers (Figure 3A), most close homologs are also homomers, while at more distant relations, many are also monomers, while heteromers are rarer. For monomers (Figure 3B), we see that at TM scores close to 1.0 (structurally identical proteins), all homologs are monomers, suggesting that our monomer filtering, as described in 2.1, worked. However, a lower TM score cut-off also includes the two other stoichiometry classes. For negative heterodimers (Figure 3C), we observe a similar trend, where heteromeric homologs are most frequent at high TM scores, while monomers become more frequent at more distant relationships.

In Figure 3D-F and supplementary Figure S2, the Stoichiometry fractions have been separated for proteins predicted with ranking confidence above or below 0.8. In Figure 3D-F, we focus on the fraction of homomultimers at different TM score cut-offs, while Figure S2 includes the fractions for all three stoichiometry categories. When we separate proteins into high- and low-confidence groups, a clear pattern emerges.

The pattern is that the fraction of homomultimers differs in the separated sets. For monomers and neg heterodimers, the samples with high scores also have a higher fraction of homomultimers in their homologs. This is especially clear for neg heterodimers. The same pattern holds for homodimers, and the fraction of homomultimers among the highly scored proteins is much higher than in the negative datasets. However, the low-scoring proteins have a slightly higher fraction than the negatives at a TM Score cut-off of 0.9, but at lower TM Score cut-offs, the fraction is similar to that of the highly scored negative datasets. This may explain why AlphaFold has a harder time with these proteins in particular, since it is difficult to distinguish positives and negatives here based on their homology information.

We find similar results when using MMseqs2 to detect sequence-based homology instead, see Figure S4 in the Supplementary material. The major difference between the results of the sequence-based and structure-based homology search methods is found in the negative heterodimers and monomers with ranking confidence above 0.8 (compare S2H-I and S4H-I). Both subsets have a higher fraction of heteromutimeric homologs for the MMseqs2 results. However, the negative heterodimers in Figure S4F and I also show a higher fraction of homomultimers for samples with a ranking confidence above 0.8, mimicking the Foldseek results.

In summary, it is clear that stoichiometry is poorly conserved across distant homologs for all protein types (monomers, heterodimers, and homodimers). At a TM score similarity of 0.9, only about 60% of the homo- and heterodimeric proteins have the same stoichiometry, whereas for monomers it is about 80%. Further, it can be noted that for incorrectly classified proteins (both low-scoring homodimers and high-scoring monomers and heterodimers), the fraction of (close) homologs with “correct” stoichiometry is lower than for correctly classified proteins. This indicates that this information can be used to improve the classification.

### Individual Performance of Features

We used three types of features, structural-, AlphaFold- and homology-based to improve the identification of homodimeric proteins, see Methods Section for details. For each feature, we plotted a precision-recall curve and found that many features in the same group behave similarly (see supplementary Figure S1 and Table S3). This is especially true for the group that contains AlphaFold confidence metrics and structural consensus. As expected, the AlphaFold confidence metrics performed well, but other features come near for some specific metrics.

For the group of features based on homology information, the higher TM score cut-offs seem to contain more valuable information, which is in line with Figure 3A. The feature *best TM Score (all)* seems to contain little information, since the AUPR is the same as a random classifier (AUPR=0.5). This feature is just the score of the highest Foldseek hit regardless of its stoichiometry, which is understandably not very useful for homodimer prediction.

The interface features inspired by SPOC (except *best pAE min*) all have AUPR between 0.75 and 0.87, suggesting performance on par with AlphaFold confidence metrics. However, the other metrics do not quite reach the performance of the homomultimer fractions or the AlphaFold confidence metrics.

### Final Clusters

As mentioned in Section 2.6 in Methods, we needed to reduce the number of features since it was infeasible to try all 44! combinations. We first clustered the 23 homology features into 6 clusters. We then chose the feature with the highest true-positive rate (TPR) at a 1% false-positive rate (FPR) from each cluster for further feature selection. In addition, the TPR at 5% FPR was used as follows: if in two clusters different members had the highest TPR at 1% and 5% FPR, both were selected for the next step, resulting in 8 features from 6 clusters. The clusters are shown in the Supplementary Material, Figure S5A. The 8 selected homology features were combined with all other features, yielding 29 features. 29 features gave us a total of 29! == 1.0888869 * 10^28^ combinations, which was still computationally infeasible. Therefore, we clustered these 29 features into 11 clusters. The dendrogram of the resulting features can be found in the Supplementary Material, Figure S5B.

### Improving True Positive Rates using Logistic Regression

We combined either 0 or 1 features from each of the 11 clusters into a logistic regression function. We also tried excluding the homology information, and in this case, we were able to test all combinations. Both functions are able to improve the TPR@1%FPR when compared with AlphaFold confidence metrics (see Table 4). The improvement is greater when including homology information: 0.42 vs 0.28 (vs 0.19 for the best AlphaFold metric). The precision-recall curves of the logistic regression models and their features are shown in Figure 4A-C. Here, it is apparent that the features of the logistic regression models vary greatly, indicating complementary behaviour. The exact features included in each model are listed in Table 3. The results of the two logistic regression models show that including homology information significantly improves TPR at 1% compared to using AlphaFold confidence metrics.

**Fig. 4.**
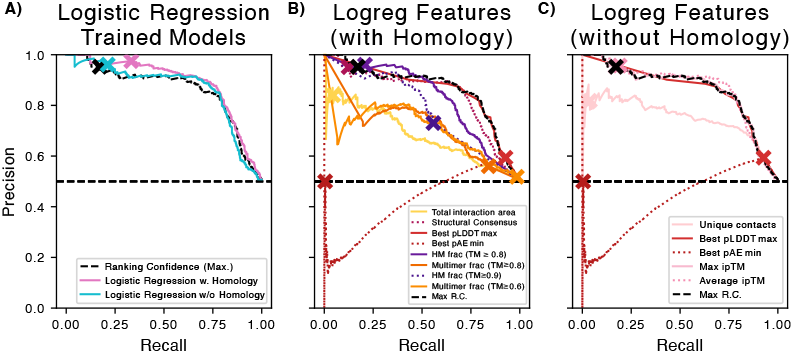
Precision-recall curves. The “X”-s mark where the FPR is 1%. A) The two logistic regression functions and the highest ranking confidence for each model. B) The precision-recall curves for the features included in the logistic regression function with homology information (Foldseek). C) The precision-recall curves for the features used for the logistic regression function without any homology features (Foldseek).

**Table 3.** Features included in the logistic regression models.

| Feature | LogReg<br>w H.I.* | LogReg<br>w/o H.I.* |
| --- | --- | --- |
| <u>Homology Information</u> |  |  |
| HM Fraction, TM Score $\geq 0.8$ | ✓ | — |
| HM Fraction, TM Score $\geq 0.9$ | ✓ | — |
| Multi Fraction, TM Score $\geq 0.6$ | ✓ | — |
| Multi Fraction, TM Score $\geq 0.8$ | ✓ | — |
| <u>Interface Information</u> |  |  |
| Total Interaction Area | ✓ | — |
| Num Unique Contacts | — | ✓ |
| Best pLDDT max | ✓ | ✓ |
| Best pAE min | ✓ | ✓ |
| <u>AF Confidence</u> |  |  |
| <u>& Structural Consensus</u> |  |  |
| Max ipTM | — | ✓ |
| Average ipTM | — | ✓ |
| Structural Consensus | ✓ | — |

\*H.I. = Homology Information.

**Table 4.**
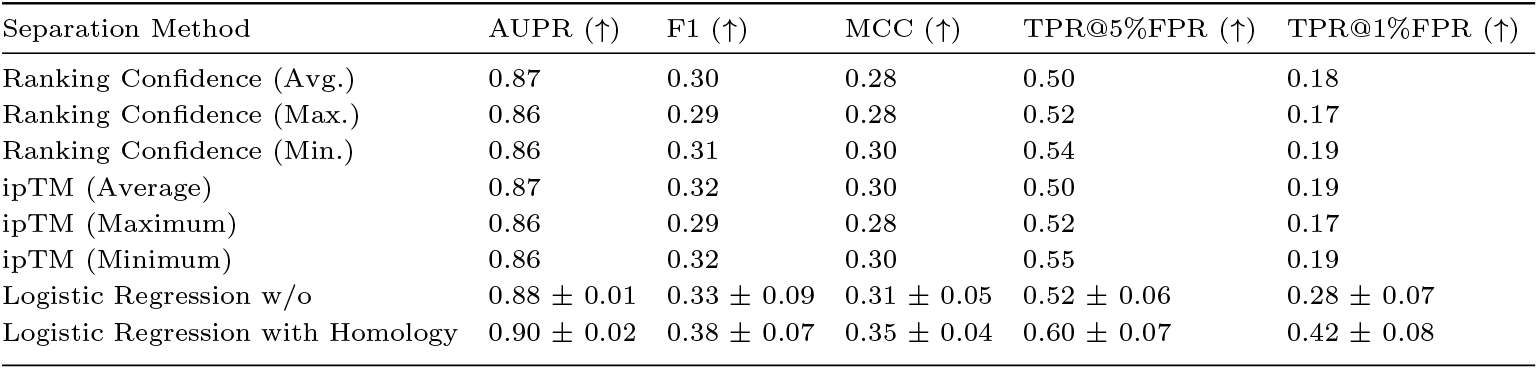
Area under the precision-recall curve (AUPR), false positive rate (FPR), false negative rate (FNR), F1-score and MCC-score for AlphaFold vs Logistic Regression classifier. All values are calculated using the highest-ranking confidence for each pdbid and the threshold used, which yields a FPR as close to 1% as possible. The logistic regression results are from a 5-fold cross-validation scheme with the standard error of the mean shown.

| Separation Method | AUPR (↑) | F1 (↑) | MCC (↑) | TPR@5%FPR (↑) | TPR@1%FPR (↑) |
| --- | --- | --- | --- | --- | --- |
| Ranking Confidence (Avg.) | 0.87 | 0.30 | 0.28 | 0.50 | 0.18 |
| Ranking Confidence (Max.) | 0.86 | 0.29 | 0.28 | 0.52 | 0.17 |
| Ranking Confidence (Min.) | 0.86 | 0.31 | 0.30 | 0.54 | 0.19 |
| ipTM (Average) | 0.87 | 0.32 | 0.30 | 0.50 | 0.19 |
| ipTM (Maximum) | 0.86 | 0.29 | 0.28 | 0.52 | 0.17 |
| ipTM (Minimum) | 0.86 | 0.32 | 0.30 | 0.55 | 0.19 |
| Logistic Regression w/o | 0.88 ± 0.01 | 0.33 ± 0.09 | 0.31 ± 0.05 | 0.52 ± 0.06 | 0.28 ± 0.07 |
| Logistic Regression with Homology | 0.90 ± 0.02 | 0.38 ± 0.07 | 0.35 ± 0.04 | 0.60 ± 0.07 | 0.42 ± 0.08 |

This can be explained by the individual performance of these metrics. When homology information features are combined with features from other groups, we can improve performance beyond that of either group alone. However, when we remove the homology information from the logistic regression function, we can only slightly improve the TPR. This fact clearly shows the importance of the homology information features.

Interestingly, both logistic regression models include features that perform poorly individually, such as *Best pAE min*. This shows the power of combining features and suggests that the combined features compensate for each other’s weaknesses. In this study, the aim was to use a simple model; however, a more advanced model, in conjunction with additional data and these “optimal” features, could potentially improve the TPR at 1% FPR further.

## Conclusions

In this study, we assessed AlphFold’s ability to identify proteins that form homodimers. First, we surprisingly found that AlphaFold produced a higher fraction of false positives for homodimers, i.e., proteins that are either monomers or part of a heterodimeric protein pair, when predicted as homodimers. This is in stark contrast to the well-studied ability of AlphaFold to identify heterodimeric interactions. We found that a possible reason is that many homologs of these high-scoring non-homodimeric proteins form homomultimers (not only homodimers). This means that “stoichiometry” is not well conserved even for quite close homologs. Next, to reduce false positives, we introduce homology-, structural, and AlphaFold-based features into a logistic regression model and find that using only 8 features is sufficient to improve homodimer prediction. Here, the most important feature is to combine the scores from AlphaFold with homology information. Using this logistic regression function, we can identify 42% of the homodimers at 1% FPR, up from only 19%. However, as the TPR for heterodimers is 63%, there is still plenty of room for improvement in accurately predicting homodimers.

## Code and Data Availability

All code and results are available on Github: https://github.com/SarahND97/alphafold-homodimers and the predicted models, fasta-files and msas are found on Zenodo, DOI: 10.5281/zenodo.17738668.

## Author contributions statement

SND and AE conceptualised the study; SND generated models, wrote scripts for analysing the results, and conducted the initial analysis. SND and AE analysed the results. SND wrote the first draft of the manuscript and made the figures. SND and AE edited and reviewed the manuscript. AE acquired the funding. AE supervised and handled all correspondence.

## Acknowledgments

The authors thank anonymous reviewers for their valuable suggestions. AE was funded by Vetenskapsrådet, Grant No. 2021-03979, and the Knut and Alice Wallenberg Foundation, Grant No. 2022.0032. The computations and data handling were enabled by the supercomputing resource Berzelius, provided by the National Supercomputer Centre at Linköping University, the Knut and Alice Wallenberg Foundation, and SNIC, grant numbers SNIC 2021/5-297 and Berzelius-2021-29. The authors also express their gratitude to the rest of the Arne Elofsson lab for valuable discussions and input.

## Supplementary Material

**Fig. S1.**
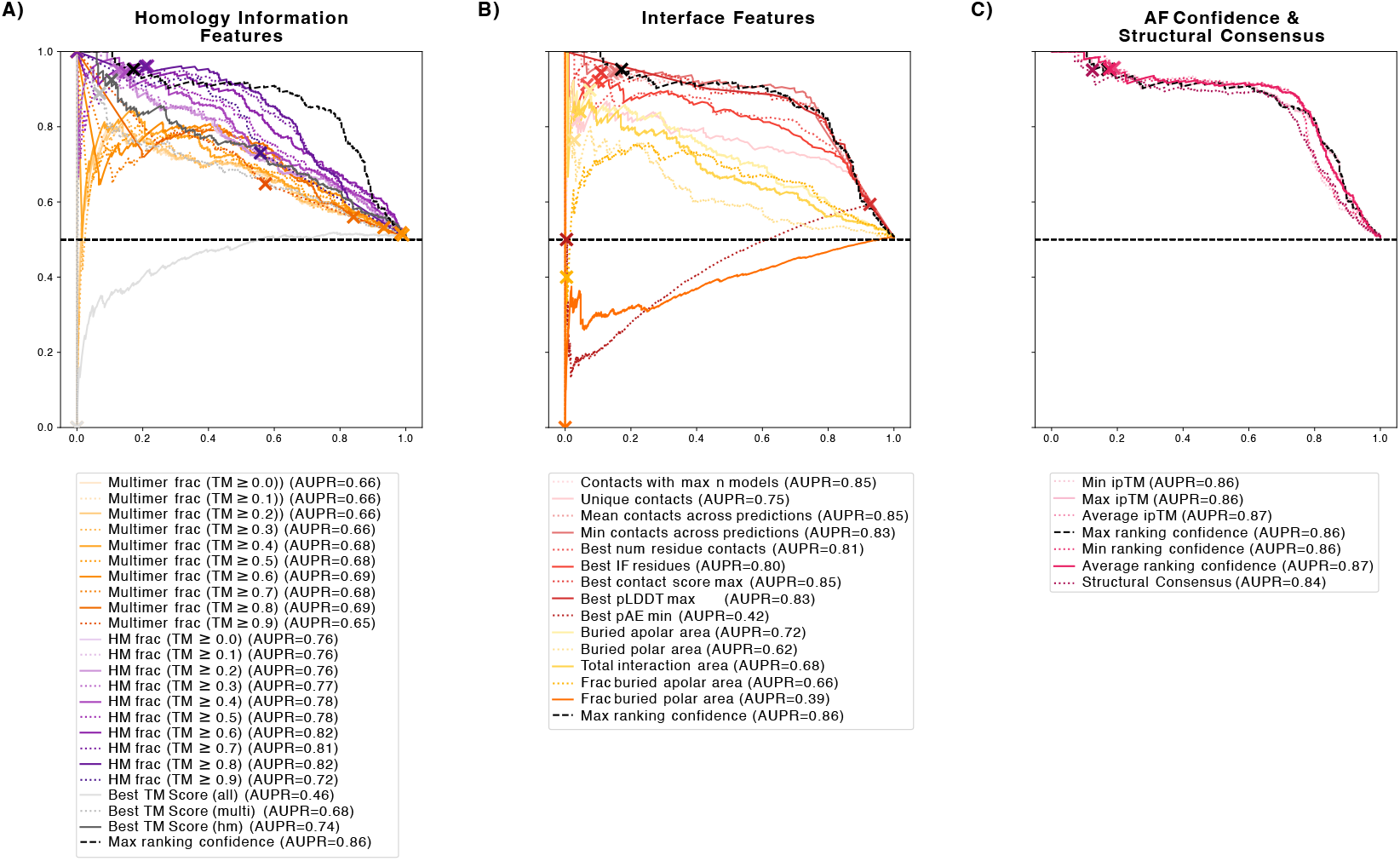
Precision-recall curve for the three groups of features that we tried out. The cross represents where the FPR is approximately 1%. The maximum ranking confidence is seen in all three figures as a reference. A) The homology information features drawn from the Foldseek matches. B) The interface features from SPOC and FreeSASA. C) The AlphaFold (AF) confidence features and structural consensus, comparison of the predicted structures to each other.

**Fig. S2.**
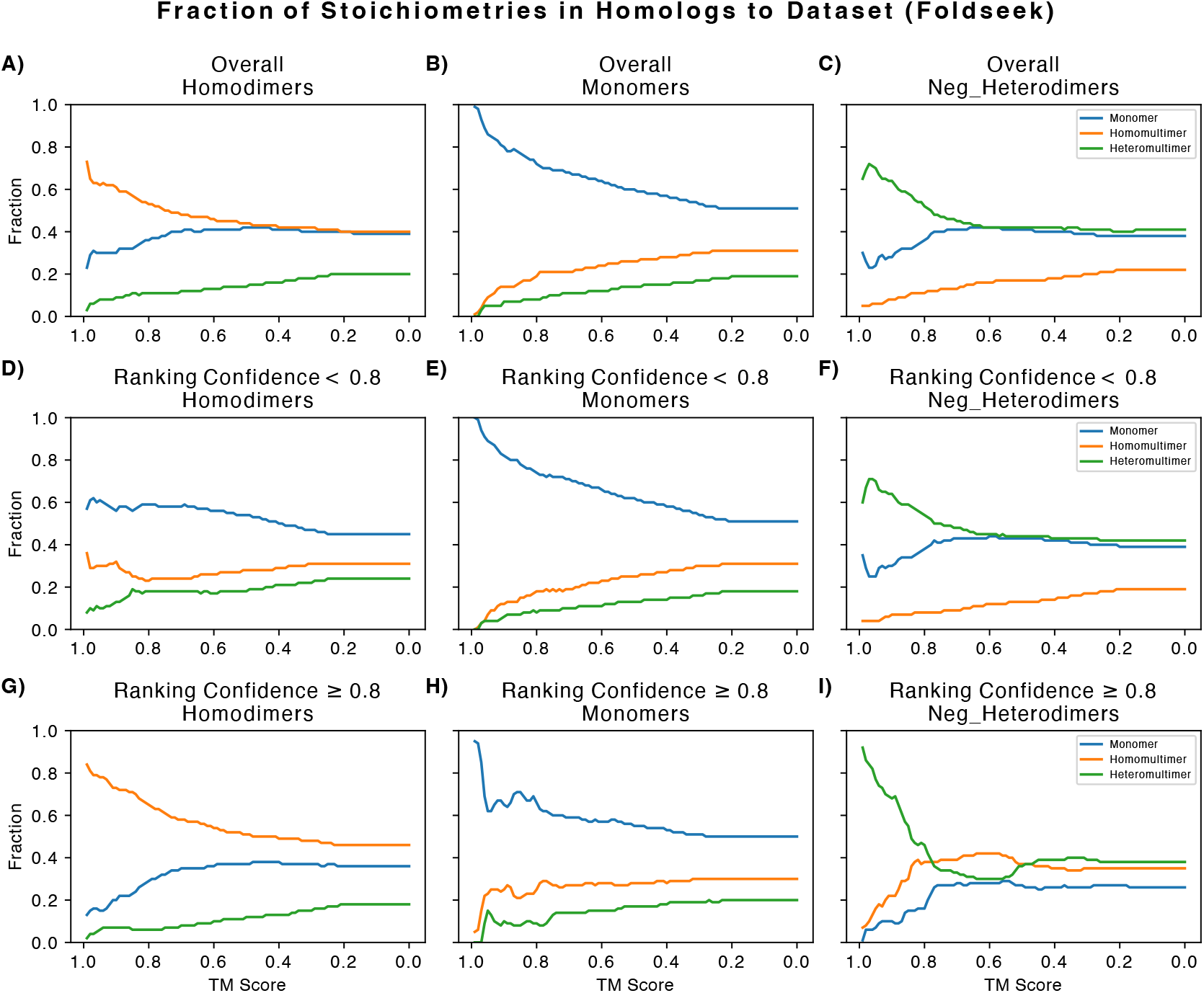
All categories of stoichiometry fractions split by ranking confidence == 0.8. In main text only homomultimeric fractions were included.

**Fig. S3.**
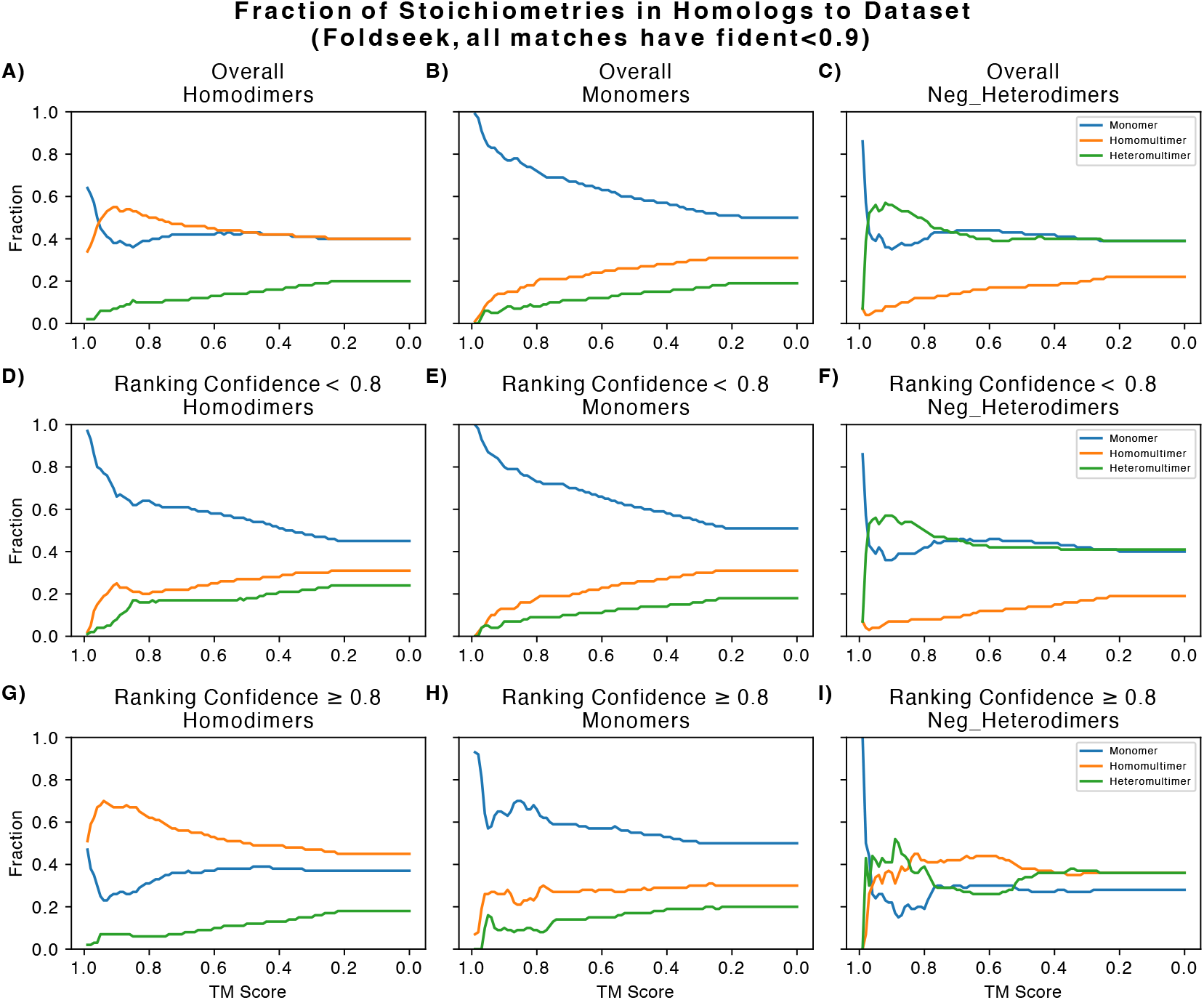
Mutimeric fractions split by ranking confidence. In this figure the matches with more than 90% sequence identity to the query were filtered out.

**Fig. S4.**
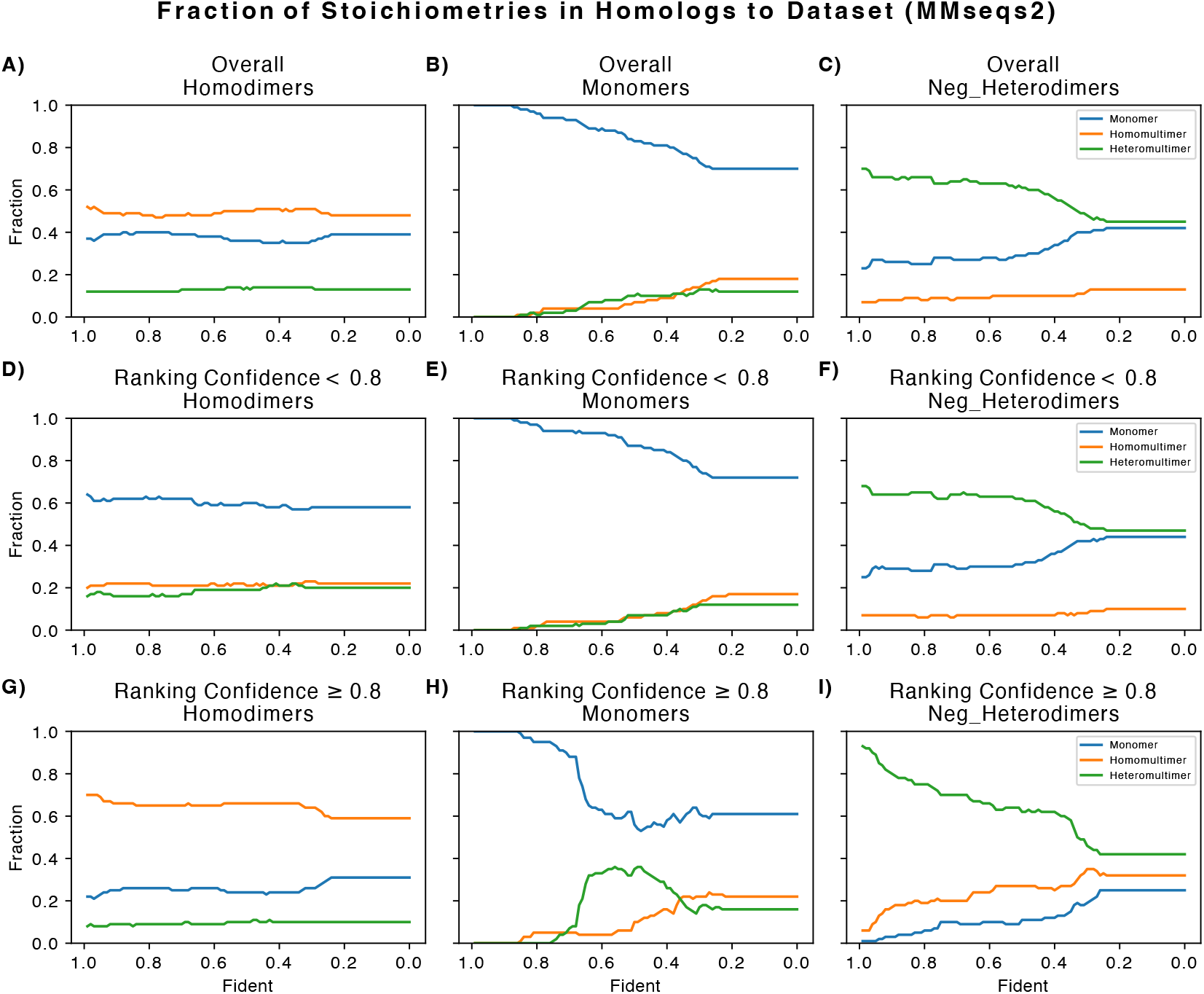
In this figure homologs were found on sequence level using MMseqs2 instead of on the structural level. Instead of using TM Score as cut-offs we used Fident. We can see a similar pattern to the structural homologs with a few exceptions.

**Fig. S5.**
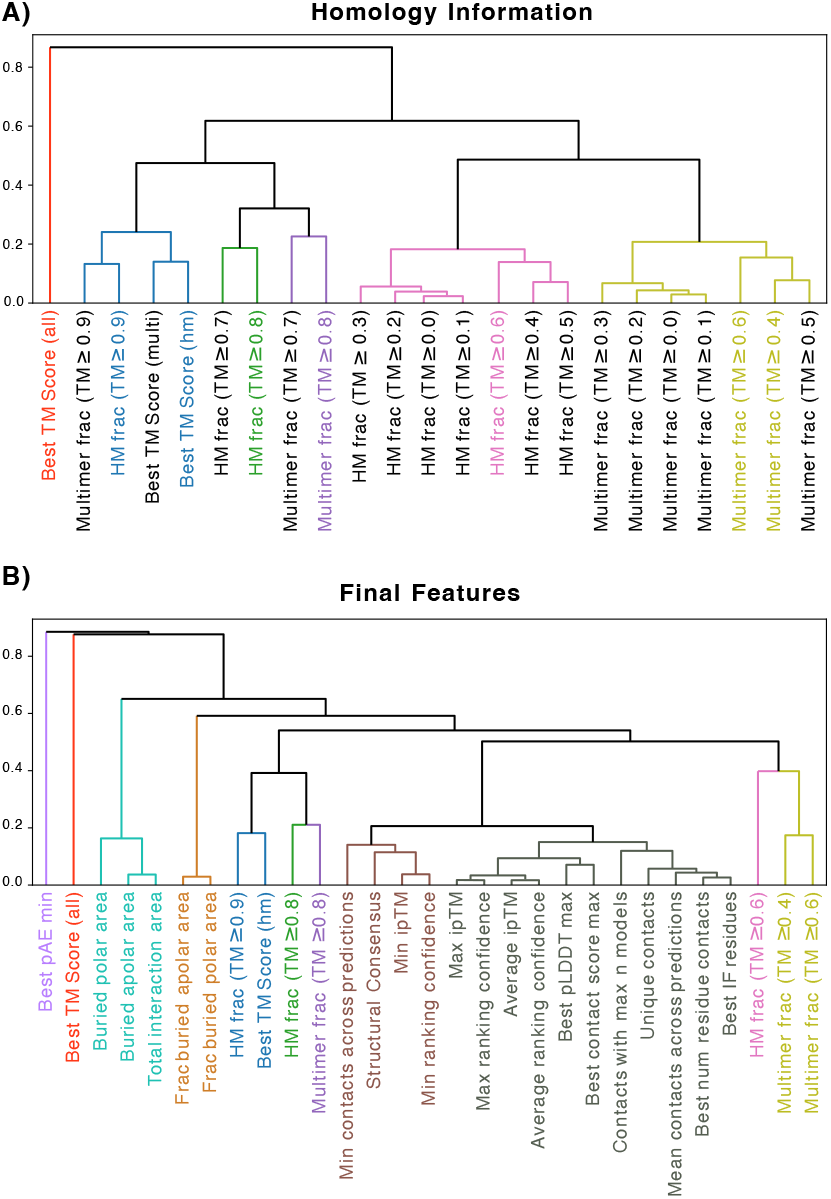
Dendrograms showing the final clustering A) Dendrogram showing the resulting clusters after the hierarchical clustering of the 23 homology features. The 8 feature names in colour are the ones chosen to be included in B). B) The final feature clustering. Note that the correlation values are altered slightly for easier visualisation.

**Table S1.** Results on the homodimeric dataset using different AlphaFold version and QSProteome from (13). The highest score for each metric is marked in bold.

| Feature | AUPR (↑) | F1@0.8 (↑) | MCC@0.8 (↑) | TPR@1%FPR (↑) | TPR@5%FPR (↑) |
| --- | --- | --- | --- | --- | --- |
| AF2.3 Max ranking confidence | 0.86289 | **0.79038** | 0.60853 | 0.17167 | 0.51931 |
| AF2.3 Min ranking confidence | 0.85943 | 0.73913 | 0.59092 | 0.18884 | 0.53863 |
| AF2.3 Average ranking confidence | 0.86886 | 0.76580 | 0.61405 | 0.17597 | 0.49571 |
| AF2.3 Min ipTM | 0.85819 | 0.73359 | 0.58645 | 0.19099 | 0.54721 |
| AF2.3 Max ipTM | 0.86370 | 0.78893 | 0.60967 | 0.16953 | 0.51502 |
| AF2.3 Average ipTM | 0.86908 | 0.76522 | **0.61485** | 0.19099 | 0.50000 |
| AF3 Max ranking score | 0.90089 | 0.75510 | 0.54978 | 0.13845 | 0.55515 |
| AF3 Min ranking score | 0.90295 | 0.70095 | 0.51274 | 0.12730 | 0.56738 |
| AF3 Avg ranking score | **0.90664** | 0.71979 | 0.52634 | 0.13279 | 0.57296 |
| AF3 Avg ipTM | 0.90413 | 0.71141 | 0.52335 | 0.18240 | 0.58240 |
| AF3 Min ipTM | 0.90132 | 0.68226 | 0.49742 | 0.14833 | 0.57339 |
| AF3 Max ipTM | 0.90059 | 0.73438 | 0.53118 | 0.20043 | 0.57926 |
| ColabFold (AF2.0) Min ipTM* | 0.86018 | 0.69693 | 0.55513 | 0.21212 | 0.58970 |
| ColabFold (AF2.0) Average ipTM* | 0.86345 | 0.74359 | 0.60131 | 0.20139 | **0.60880** |
| ColabFold (AF2.0) Max ipTM* | 0.85128 | 0.76457 | 0.57030 | 0.19187 | 0.49596 |
| QSProteome Max PAE4Con3 | 0.86050 | 0.59854 | 0.48076 | **0.21674** | 0.53004 |

\*Original AlphaFold2.0 did not include an ipTM score. This is a ColabFold implementation.

**Table S2.**
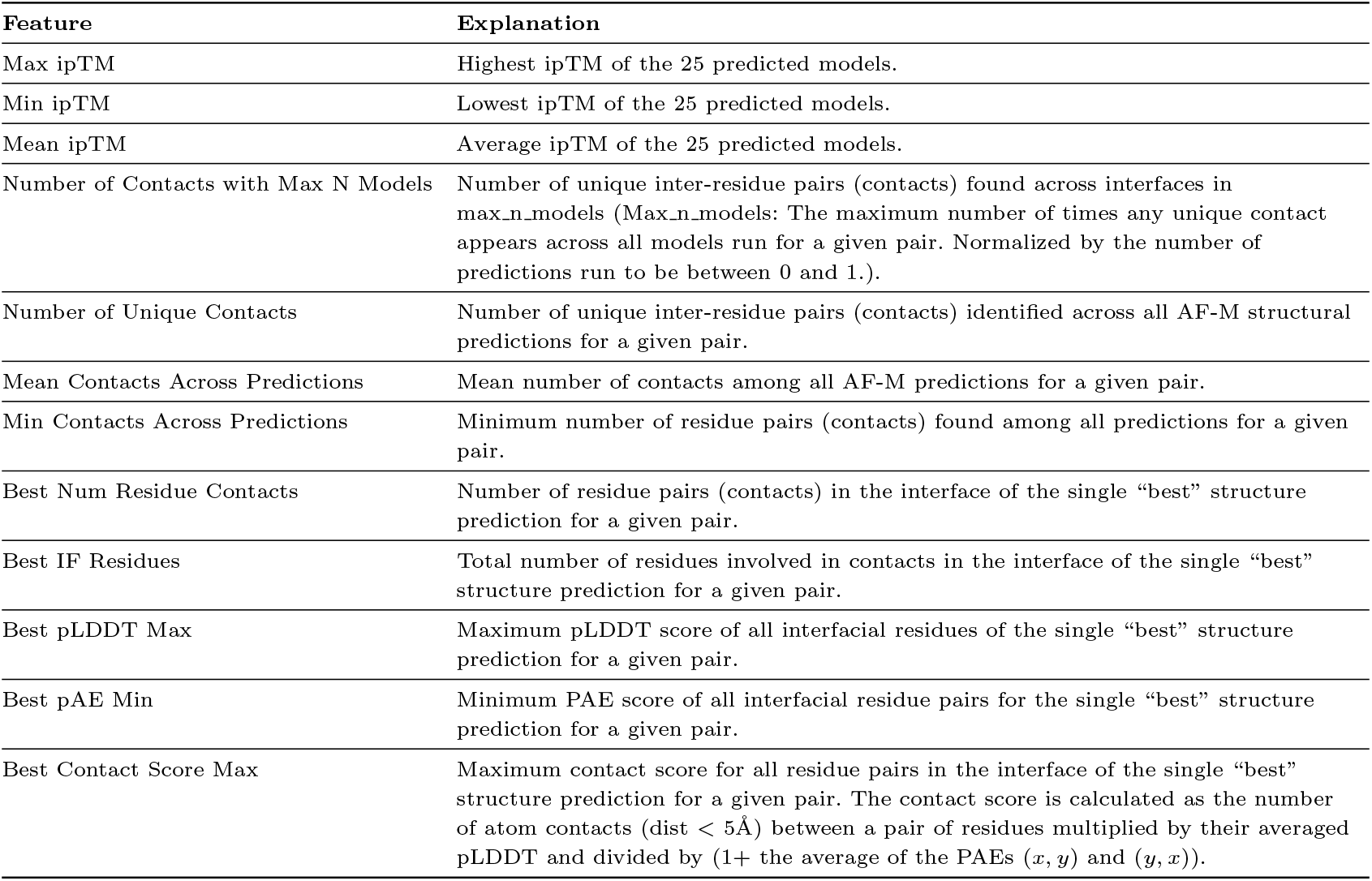
Explanation of each feature retrieved using code from SPOC. Taken from Table S3 of the supplementary material of Predictomes (18).

| Feature | Explanation |
| --- | --- |
| Max ipTM | Highest ipTM of the 25 predicted models. |
| Min ipTM | Lowest ipTM of the 25 predicted models. |
| Mean ipTM | Average ipTM of the 25 predicted models. |
| Number of Contacts with Max N Models | Number of unique inter-residue pairs (contacts) found across interfaces in max_n_models (Max_n_models: The maximum number of times any unique contact appears across all models run for a given pair. Normalized by the number of predictions run to be between 0 and 1.). |
| Number of Unique Contacts | Number of unique inter-residue pairs (contacts) identified across all AF-M structural predictions for a given pair. |
| Mean Contacts Across Predictions | Mean number of contacts among all AF-M predictions for a given pair. |
| Min Contacts Across Predictions | Minimum number of residue pairs (contacts) found among all predictions for a given pair. |
| Best Num Residue Contacts | Number of residue pairs (contacts) in the interface of the single “best” structure prediction for a given pair. |
| Best IF Residues | Total number of residues involved in contacts in the interface of the single “best” structure prediction for a given pair. |
| Best pLDDT Max | Maximum pLDDT score of all interfacial residues of the single “best” structure prediction for a given pair. |
| Best pAE Min | Minimum PAE score of all interfacial residue pairs for the single “best” structure prediction for a given pair. |
| Best Contact Score Max | Maximum contact score for all residue pairs in the interface of the single “best” structure prediction for a given pair. The contact score is calculated as the number of atom contacts ($\text{dist} < 5\text{\AA}$) between a pair of residues multiplied by their averaged pLDDT and divided by $(1 + \text{the average of the PAEs } (x, y) \text{ and } (y, x))$. |

**Table S3.** The results of individual performance of features collected for homodimer performance improvement.

| Feature | AUPR (↑) | F1@1%FPR (↑) | MCC@1%FPR (↑) | TPR@1%FPR (↑) | TPR@5%FPR (↑) |
| --- | --- | --- | --- | --- | --- |
| <u>Homology Features</u> |  |  |  |  |  |
| <u>Sequence identity $\leq 0.5$</u> |  |  |  |  |  |
| Multimer frac (TM $\geq 0.0$) | 0.66 | 0.00 | -0.07 | 0.00 | 0.15 |
| Multimer frac (TM $\geq 0.1$) | 0.66 | 0.00 | -0.07 | 0.00 | 0.15 |
| Multimer frac (TM $\geq 0.2$) | 0.66 | 0.00 | -0.07 | 0.00 | 0.15 |
| Multimer frac (TM $\geq 0.3$) | 0.65 | 0.00 | -0.07 | 0.00 | 0.17 |
| Multimer frac (TM $\geq 0.4$) | 0.68 | 0.03 | 0.01 | 0.01 | 0.21 |
| Multimer frac (TM $\geq 0.5$) | 0.68 | 0.00 | 0.00 | 0.02 | 0.19 |
| Multimer frac (TM $\geq 0.6$) | 0.71 | 0.00 | 0.00 | 0.02 | 0.13 |
| Multimer frac (TM $\geq 0.7$) | 0.70 | 0.00 | 0.00 | 0.02 | 0.10 |
| Multimer frac (TM $\geq 0.8$) | 0.72 | 0.00 | 0.00 | 0.03 | 0.14 |
| Multimer frac (TM $\geq 0.9$) | 0.71 | 0.00 | 0.00 | 0.03 | 0.16 |
| HM frac (TM $\geq 0.0$) | 0.76 | 0.25 | 0.25 | 0.14 | 0.27 |
| HM frac (TM $\geq 0.1$) | 0.76 | 0.25 | 0.25 | 0.14 | 0.27 |
| HM frac (TM $\geq 0.2$) | 0.76 | 0.24 | 0.24 | 0.14 | 0.27 |
| HM frac (TM $\geq 0.3$) | 0.77 | 0.23 | 0.23 | 0.13 | 0.29 |
| HM frac (TM $\geq 0.4$) | 0.78 | 0.26 | 0.25 | 0.15 | 0.34 |
| HM frac (TM $\geq 0.5$) | 0.79 | 0.30 | 0.28 | 0.18 | 0.37 |
| HM frac (TM $\geq 0.6$) | 0.82 | **0.35** | **0.32** | **0.21** | 0.43 |
| HM frac (TM $\geq 0.7$) | 0.82 | 0.12 | 0.14 | 0.06 | 0.48 |
| HM frac (TM $\geq 0.8$) | 0.83 | 0.34 | 0.31 | **0.21** | 0.50 |
| HM frac (TM $\geq 0.9$) | 0.78 | 0.00 | 0.00 | 0.09 | 0.40 |
| Best TM Score (all) | 0.50 | 0.00 | -0.07 | 0.00 | 0.02 |
| Best TM Score (multi) | 0.68 | 0.13 | 0.15 | 0.07 | 0.16 |
| Best TM Score (hm) | 0.74 | 0.19 | 0.20 | 0.11 | 0.27 |
| <u>Interface Features</u> |  |  |  |  |  |
| Contacts with max n models | 0.86 | 0.26 | 0.25 | 0.15 | 0.45 |
| Unique contacts | 0.75 | 0.07 | 0.09 | 0.04 | 0.22 |
| Mean contacts across predictions | 0.86 | 0.25 | 0.25 | 0.14 | 0.43 |
| Min contacts across predictions | **0.87** | 0.27 | 0.26 | 0.15 | 0.50 |
| Best num residue contacts | 0.81 | 0.14 | 0.16 | 0.08 | 0.36 |
| Best IF residues | 0.81 | 0.18 | 0.19 | 0.10 | 0.36 |
| Best contact score max | 0.85 | 0.24 | 0.24 | 0.14 | 0.44 |
| Best pLDDT max | 0.86 | 0.00 | 0.00 | 0.09 | 0.44 |
| Best pAE min | 0.52 | 0.00 | 0.00 | 0.002 | 0.01 |
| Buried apolar area | 0.71 | 0.14 | 0.16 | 0.08 | 0.22 |
| Buried polar area | 0.62 | 0.05 | 0.06 | 0.03 | 0.14 |
| Total interaction area | 0.68 | 0.09 | 0.10 | 0.05 | 0.19 |
| Frac buried apolar area | 0.66 | 0.01 | -0.02 | 0.01 | 0.12 |
| Frac buried polar area | 0.66 | 0.01 | -0.02 | 0.01 | 0.12 |
| <u>AF Confidence Metrics</u> |  |  |  |  |  |
| <u>& Structural Consensus</u> |  |  |  |  |  |
| Min ipTM | 0.86 | 0.32 | 0.30 | 0.19 | **0.55** |
| Max ipTM | 0.86 | 0.29 | 0.28 | 0.17 | 0.52 |
| Average ipTM | **0.87** | 0.32 | 0.30 | 0.19 | 0.50 |
| Max ranking confidence | 0.86 | 0.29 | 0.28 | 0.17 | 0.52 |
| Min ranking confidence | 0.86 | 0.31 | 0.30 | 0.19 | 0.54 |
| Average ranking confidence | **0.87** | 0.30 | 0.28 | 0.18 | 0.50 |
| Structural Consensus | 0.84 | 0.23 | 0.23 | 0.13 | 0.42 |

